# Inference of evolutionary jumps in large phylogenies using Lévy processes

**DOI:** 10.1101/089276

**Authors:** Pablo Duchen, Christoph Leuenberger, Sándor M. Szilágyi, Luke Harmon, Jonathan Eastman, Manuel Schweizer, Daniel Wegmann

## Abstract

While it is now widely accepted that the rate of phenotypic evolution may not necessarily be constant across large phylogenies, the frequency and phylogenetic position of periods of rapid evolution remain unclear. In his highly influential view of evolution, G. G. Simpson supposed that such evolutionary jumps occur when organisms transition into so called new adaptive zones, for instance after dispersal into a new geographic area, after rapid climatic changes, or following the appearance of an evolutionary novelty. Only recently, large, accurate and well calibrated phylogenies have become available that allow testing this hypothesis directly, yet inferring evolutionary jumps remains computationally very challenging. Here, we develop a computationally highly efficient algorithm to accurately infer the rate and strength of evolutionary jumps as well as their phylogenetic location. Following previous work we model evolutionary jumps as a compound process, but introduce a novel approach to sample jump configurations that does not require matrix inversions and thus naturally scales to large trees. We then make use of this development to infer evolutionary jumps in *Anolis* lizards and Loriini parrots where we find strong signal for such jumps at the basis of clades that transitioned into new adaptive zones, just as postulated by Simpson's hypothesis.

---

A key goal of evolutionary biology is to understand the mechanisms by which the large phenotypic diversity seen today evolved. Our understanding of these mechanisms is improving rapidly with the advent of increasingly powerful sequencing approaches. For instance, the huge amount of molecular data has led to the resolution of phylogenetic trees encompassing entire orders. Further, methods to reliably identify substitutions that likely resulted from selection, and to accurately place them on a phylogeny have been developed. In contrast, methods to infer events of rapid evolution from phenotypic data have lagged and are mostly restricted to inferring independent evolutionary rates for different clades.

In general, quantitative studies of the evolution of phenotypic/quantitative traits date back to just a few decades. A first attempt was by Edwards et al. (1964) and Cavalli-Sforza and Edwards (1967), who modeled quantitative traits stochastically as “Brownian motion” (BM). However, given the current wealth of molecular data available, a more realistic goal is to only aim at inferring the rates at which quantitative traits evolve, while assuming the underlying phylogeny to be known. This has been successfully done using a BM model in multiple taxa. Freckleton et al. (2002), for instance, used a BM model on a given phylogeny to test if traits showed phylogenetic associations. More recently, Brawand et al. (2011) modeled gene expression evolution as BM and rejected evolution at a constant rate for several genes.

Several extensions to a basic BM model have been proposed. Butler and King (2004) were the first to implement Ornstein-Uhlenbeck (OU) processes with multiple evolutionary optima, as initially described by Hansen (1997), and recently used to describe the evolution of gene expression (e.g. Bedford and Hartl, 2009; Rohlfs et al., 2013). Other extentions to BM allow evolutionary rates to change over time. O’Meara et al. (2006), for instance, contrasted maximum likelihood (ML) estimates of evolutionary rates under BM and showed that major clades of angiosperms vastly differ in their rate of genome size evolution. More recently, Eastman et al. (2011) developed a Bayesian method to jointly infer evolutionary rates in different clades and found evidence for multiple rate shifts in body size evolution in emydid turtles. Shortly after, Slater et al. (2012) have introduced an extension to incompletely sampled phylogenies and trait data using Approximate Bayesian Computation. However, they found no evidence for an elevated rate of body size evolution in pinnipeds in comparison to terrestrial carnivores, despite considerable power. This suggests that the larger body size found in pinnipeds may be the result of rapid evolutionary changes early in the clade, rather than a change in the rate itself, and hence that models of occasional “evolutionary jumps” may often more accurately explain the evolution of quantitative traits.

According to Simpson (1944), such evolutionary jumps are triggered by shifts of lineages into different adaptive zones, either by dispersal into new geographic areas, the appearance of evolutionary novelties, key innovations, the extinction of lineages leaving niches empty, or by rapid changes in the environment (climatic or ecological). Additionally, the existence of “ecological opportunities” (Losos, 2010) might also trigger such jumps. While OU processes have been proposed to model the dynamics of adaptive landscapes (e.g. Ingram and Mahler, 2013; Uyeda and Harmon, 2014), a promising alternative is to model this type of evolution as a compound process (or L´evy process) consisting of a continuous background process and a discrete jump process. The first implementation of such a model assumed that jumps only occured at speciation events (Bokma, 2008), but Landis et al. (2013) recently described L´evy processes in a much more general way and showed that while the likelihood functions of most of these models are intractable, inference is possible under a Bayesian framework. For instance, when modeling the evolution of quantitative traits as a Poisson compound process, in which traits are assumed to evolve under BM with occasional jumps that occur as a Poisson process on the tree, the likelihood can be calculated analytically when conditioning on a jump configuration (a placement of jumps on the tree). Under the assumption that jump effects are normally distributed, a jump configuration can be seen as simply stretching the branches of the tree on which they occur, and the likelihood is then given by a multivariate normal distribution with the variance-covariance matrix resulting from the stretched tree. The numerical integration is then limited to sampling jump configurations, which is readily done using Markov Chain Monte Carlo (MCMC).

Unfortunately, two computational challenges prohibit the application of this approach to larger trees. First, the space of jump configurations grows exponentially with tree size, leading to very long MCMC chains. Second, the evaluation of the likelihood requires the computation of the inverse of the variance-covariance matrix, which is computationally very demanding since it scales exponentially with tree size (Tung Ho and Ané, 2014). Here, we address these computational issues using an empirical Bayes approach in which we first infer the hierarchical parameters of the Brownian and Poissonian processes using Maximum Likelihood, and then fix those when inferring posterior probabilities on jump locations. This approach allows us to run MCMC chains with fixed hierarchical parameters, for which we find a computationally highly efficient approach that does not require matrix inversions. As a result, this approach readily scales up to very large phylogenies.

We then demonstrate the power and accuracy of our approach with extensive simulations and find that our approach hardly misses any jumps with a meaningful strength. We then illustrate the usefulness of our approach by identifying evolutionary jumps in *Anolis* lizards and Loriini parrots, two well-studied groups for which morphological data is available. We identify few but important evolutionary jumps in both groups, suggesting such periods of rapid evolutionary change to be rare but crucial in shaping the morphological diversity observed today.

## THEORY

### The null hypothesis: Brownian motion

We first consider a Brownian motion (BM) process on a phylogenetic tree 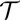 with root 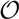 where time is measured in the unit of the branch lengths. The process starts at 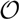 with value *μ* ∈ ℝ (root state) and then proceeds with variance 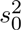 along the branches. The values of the BM process, as observed at the *L* leaves, give rise to the random vector

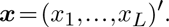

Let us fix the notation: The lenghts of the (inner and outer) branches of 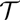 are called *τ*_1_,…,*τ_B_* where *B* is the number of branches. For two leaves *i,j* we denote by ***T***_0_ = (*τ_ij_*) the length of their common branch in 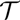 as measured from the root 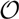. Now, under the assumption of a pure BM, and defining **1** = (1,1,…,1)′, the values ***x*** at the leaves have the multivariate normal distribution

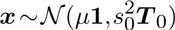

or written more conveniently:

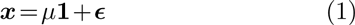

with 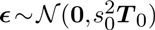. Since ***T***_0_ is positive definite and symmetric, it has a symmetric and positive definite square root ***Q***, i.e. ***Q***^2^ = ***T***_0_. Multiplying both sides of (1) with ***Q***^*−1*^ we get the homoskedastic model

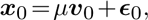

where ***x***_0_ = ***Q***^−1^***x***, ***v***_0_ = ***Q***^−1^**1**, and 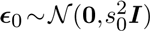. For this we have the usual OLS estimators (see e.g. Davidson and MacKinnon (2004), ch. 3.2)

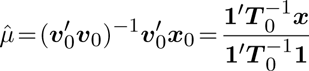

and

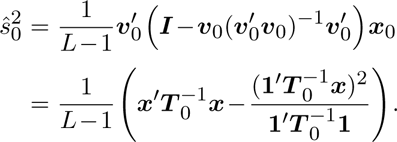

### Lévy process

We now extend the BM model by super-imposing an independent Poissonian jump-process with rate *λ*. The jumps shall be normally distributed with zero mean and variance 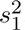. The (unobservable) random vector

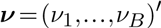

counts the number of Poisson events (jumps) on each of the *B* branches. By assumption,

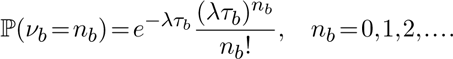

For a multi-index ***n*** = (*n*_1_,…,*n_B_*), we have

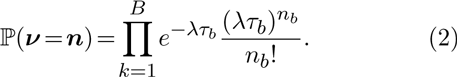

Recall that for two leaves *i,j* we denote by *τ_ij_* the length of their common branch in 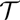 as measured from the root 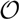. In particular, *τ_ii_* is the distance (sum of branch lengths) of the leaf *i* from the root 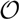.

We denote by *n_ij_* for two leaves *i,j* the number of Poisson events along the common branch of length *τ_ij_*. Conditional on ***ν*** = ***n*** = (*n*_1_,…,*n_B_*), the random vector ***x*** is multivariate normal with mean *μ***1** and the *L* × *L* variance-covariance matrix **Σ**(***n***) = (*σ_ij_*(***n***)) where

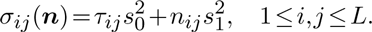

The conditional density of ***x*** given ***ν*** = ***n*** is

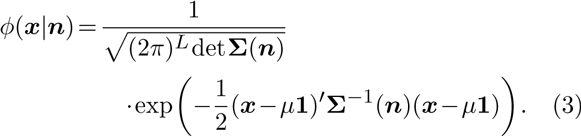

The likelihood of ***x*** given the four parameters *μ* (root state), *s*_0_ (Brownian motion) and *λ*,*s*_1_ (Poissonian jump process) is the mixture distribution

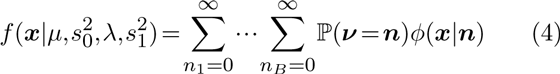

where we used expressions (2) and (3). It is not hard to show that

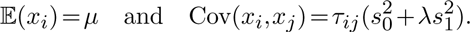

### Inference under the Lévy process

Here we develop a computationally efficient approach to maximize the likelihood function given in equation (4). While the infinite sums in (4) prohibit an analytical solution, they are readily evaluated using numerical approaches. Landis et al. (2013), for instance, proposed to use an MCMC approach to integrate over jump configurations. Unfortunately, however, such a solution does not scale to large trees, because the calculation of the conditional density values in (3) involves the computation of the inverse of **Σ**(***n***) and its determinant, which are computationally very demanding.

We propose to address this problem by introducing an algorithm to calculate these matrix inversions efficiently under this model. While this algorithm can readily be incorporated into the MCMC approach proposed by Landis et al. (2013), we will then propose an alternative hierarchical Bayes approach that makes even better use of it and leads to a computationally highly efficient inference approach to obtain point estimates of the parameters 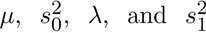, as well as posterior probabilities on the location of evolutionary jumps.

### Efficient calculation of inverses and determinants

For a symmetric non-singular matrix ***A*** and a (column) vector ***a***, we have

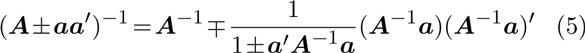

(see Izenman (2008), p. 47) and

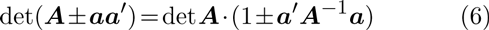

(see Anderson (2003), Corollary A.3.1). These formulae have recently been shown to speed up the calculation of the likelihodo function under Brownian motion models (Tung Ho and Ané, 2014). Here we use them to develop a fast algorithm applicable to Lévy processes.

Let us first fix some notation: For each branch *b*, we define the *L* × *L* incidence matrix 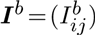 by setting 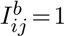 if the branch *b* is common to the pair of leaves *i,j*, and 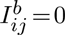 otherwise. Clearly,

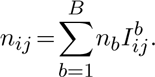

In the following we replace the parameter 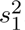 with the positive factor *α* given by

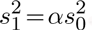

Observe that

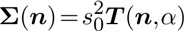

and

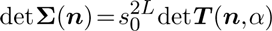

where

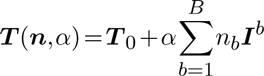

and ***T***_0_ = (*τ_ij_*). Finally, we introduce for *b* = 1,…,*B* the (column) vectors ***u***^*b*^, each one with *L* components. The *i*-th component 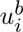 is equal to 1 if leaf *i* is subordinate to branch *b* (i.e. the path from the root 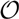 to node *i* contains branch *b*). Otherwise, if leaf *i* is not subordinate to branch *b*, then 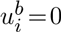. It is easy to see that ***I***^*b*^ = ***u***^*b*^(***u***^*b*^)′ and thus

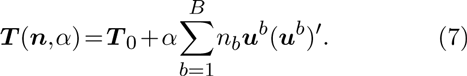

We can now apply formulae (5) and (6) to obtain the following iterative scheme for the computation of ***T***^−1^(***n***,*α*) and det ***T***(***n***,*α*):

First, determine 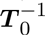 and det ***T***_0_. Then, for each term with *n_b_* > 0 in the sum (7), update ***T***_*b* − 1_ to ***T***_*b*_ etc. as follows: Let

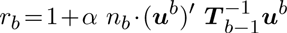

and calculate

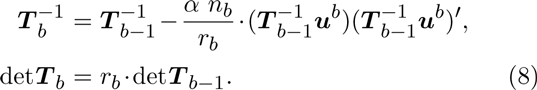

When all non-zero terms in (7) have been considered, we arrive at 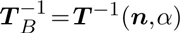 and det ***T****B* = det ***T***(***n***,*α*). Observe that in this scheme, the only matrix inverse that ever has to be determined is 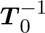. The number of non-zero *n_b_* will frequently be small compared to *B* and so will be the number of iterations (8).

*Monte Carlo EM algorithm* The scheme to calculate the inverse of **Σ**(***n***) allows to find the ML estimates of the parameters 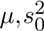 and *λ* by means of a Monte Carlo version of the classical Expectation Maximization (EM) algorithm, in which we treat the random variable = as missing (unobserved) data. While this approach does not allow us to find the ML estimate of *α*, we discuss below how this can be achieved using a simple grid search.

Recall that each iteration of the EM algorithm consists of an estimation (E) and a maximization (M) step. Let us denote the old parameters determined in the previous M-step by 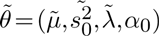, and the new parameters with respect to which the *Q*-function has to be maximized in the next M-step by 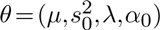, where *α*_0_ is a fixed value for *α*. The two steps of the EM algorithm are then as follows:

*Monte Carlo E-step.* Simulate stochastically *K* vectors ***n***_*k*_ according to the multi-Poisson distribution ℙ(***ν*** = ***n***|*λ̃*). Determine the weights

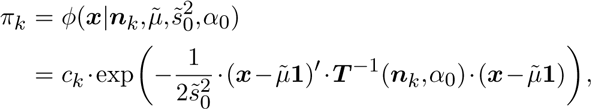

with

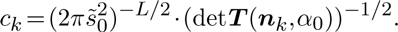

In the M-step we have to maximize the function

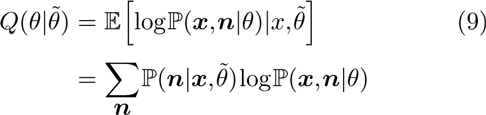

with respect to the parameters 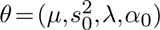 where

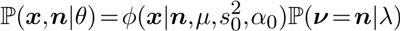

From Bayes' theorem we have

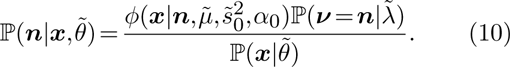

Thus, according to our Monte Carlo scheme and up to the factor 1/ℙ(***x***|*θ̃*), the infinite sum in (9) can be approximated by

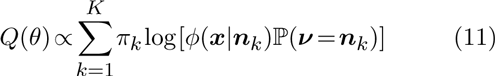

where *φ*(***x***|***n***_*k*_) and ℙ(***ν*** = ***n***_*k*_) are given by (2) and (3), respectively.

*M-step.* In this step we seek the parameters *θ̃* which maximize the sum in (11) and which will serve as “old” parameters in the next E-step. We have

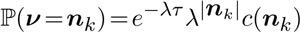

where *τ* = *τ_i_* is the total length of the tree 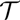, |***n***_*k*_| denotes the sum of the components of *n_k_*, and *c*(***n***_*k*_) is a factor that does not depend on any of the parameters *θ*. From this it is easy to see that

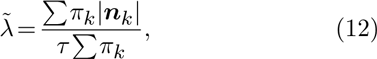

independently of the values of the other three parameters. Since we assume the value of *α* to be fixed, we can also give explicit expressions for the values of *μ* and 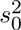 which maximize *Q*(*θ*|*α* = *α*_0_). First, determine the matrix

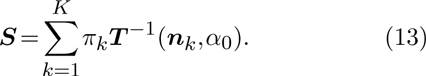

Standard calculus shows that

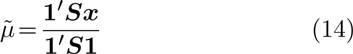

and

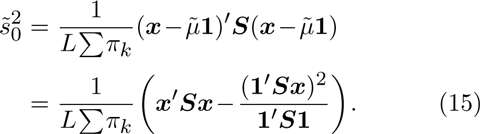

*Simulating* ***n*** *with MCMC* In this section we describe how to sample the states ***n*** from the probability distribution ℙ(***n***|***x***,*θ*) using the Metropolis scheme. (To unburden the notation in the description of the MCMC algorithm, we drop the tilde overscript on the parameters.) At each state we will need the inverse matrix ***T***^−1^ of ***T***(***n***,*α*_0_) given by (7). Start the chain e.g. at ***n*** = (0,…,0) and with 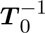.

1. Let ***n*** denote the current state of the Markov chain and ***T***^−1^ the inverse matrix of ***T***(***n***,*α*_0_). Choose an index *b* = 1,…,*B* with equal probability (or with a probability proportional to *τ_b_*) and an increment Δ*n_b_* = +1 or = −1 with probability 1/2. The candidate state ***n***′ is given by in‐ or decreasing the *b*-th index ***n*** by 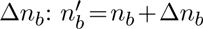.
2. Using (10) and the iteration formula (8) it is not hard to check that the Hastings ratio (proposal probability) can be calculated by

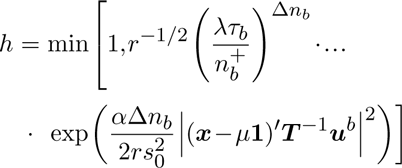

where 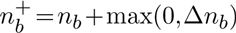 and

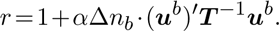

If the candidate state contains a negative component (i.e. if *n_b_* = 0 and ∆*n_b_* = − 1) then set *h* = 0. This ensures that the chain is indeed symmetric.
3. With probability *h* jump to the candidate state ***n***′, otherwise stay at ***n***. In the first case, update

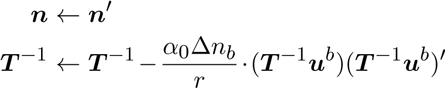

and go to step 1.

No matrix inverse must ever be calculated in this scheme thanks to the update in step 3. (To counterbalance the accumulation of numerical errors it might however be wise to occasionally calculate ***T***^−1^ = ***T***^−1^(***n***,*α*_0_) from scratch.)

After the burn-in phase, a fraction ***n***_1_,…,***n***_*M*_ of the simulated states will be retained (“thinning out”). These will be used to replace the matrix (13) in the M-step of the EM algorithm by

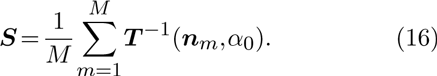

*Estimating factor α* The Monte Carlo EM algorithm proposed above, while computationally highly efficient, does not allow for the estimation of the factor *α*. We thus use a numerical approach to iteratively approach the ML estimate of *α*. Specifically, we start at a value *α*_0_ and then iteratively increase that value such that log_10_*α_t_ = log_10_*α*_*t*_ − 1* + Δ_*α*_ until the likelihood decreases. The algorithm then turns back by setting Δ_*α*_ ← −Δ_*α*_/*e* and proceeds again until the likelihood decreases. With every switch, the step size gets smaller and the estimate closer to the true MLE value. In each step we use the Monte Carlo EM algorithm described above to calculate the likelihood at the MLE estimates of all other parameters conditioned on that *α* value. In all application we set *α*_0_ = 0.1 and the initial Δ_*α*_ = 0.1 and found estimates to be accurate within five switches.

*Identifying jump locations* To infer the location of jumps on a phylogenetic tree we implement an empirical Bayes approach. As is commonly done in such a setting, we assume the ML estimates 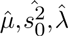 and *α̂* obtained using our Monte Carlo EM scheme are accurate and thus known constants when inferring jump locations. Under this assumption, the MCMC approach introduced above can also be used to sample configurations of jumps ***n*** from the probability distribution 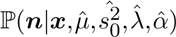. This allows us to numerically infer for each branch *k* the posterior probabilities of 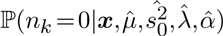 and 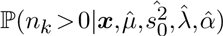, and thus to identify branches for which there is convincing evidence for an evolutionary jump.

*Implementation* We implemented the algorithm introduced here in C_++_ and optimized the code for speed. A user-friendly program to apply it to data is available at our lab website (http://www.unifr.ch/biology/research/wegmann/).

## SIMULATIONS

### Convergence

*Convergence of the MCMC* We assessed the convergence of MCMC chains by comparing parameter estimates between two independent and parallel chain runs until 10,000 jump vectors ***n*** were sampled. We run a total of 100 such chain pairs for each of two starting locations with values differing ten fold and discarded the first 100 such vectors as burn-in. We also compared two different values to thin the chains: either we sampled every 10th or every 5000th step.

Regardless of the starting values, convergence was reached rather fast but with some variation across parameters (Fig. S1). The parameter to converge fastest was *μ*, for which the difference in estimates was below 0.01 within 2,000 sampled jump vectors for 90% of all chain pairs. Similarly small differences for *s*2 and *λ* were only reached after sampling about 4,000 jump vectors (Fig. S1). Interestingly, a larger thinning did not improve convergence, suggesting that the variance in estimates is dominated by variation in the jump vectors sampled, but not by autocorrelation along the chain. For subsequent analyses, we used a thinning of 10 and sample a total of 5,000 jump vectors.

We next assessed the convergence of the MCMC for the inference of jumps on trees by assessing the difference in posterior probabilities between independent chains (Figure S2). We again run 100 chain pairs, fixed the thinning to 10 and discarded the first 100 jump vectors as burn-in. While we found convergence to be reached within less than 2,000 iterations for branches with very low (< 0.05) and very high (> 0.95) posterior probabilities, more iterations were required for branches with intermediate posterior probabilities. We found that sampling 5,000 jump vectors gave very consistent results also for inferring the location of jumps.

*Convergence of the EM for parameter inference* To test if the stochastic EM algorithm converges with the MCMC settings found above (5,000 jump vectors, a burn-in of 100 such vectors and a thinning of 10), we run the EM for a wide range of parameter values for up to 100 iterations. Since the EM algorithm is stochastic, it does not converge onto a single value unless an infinitely large sample of ***n*** vectors are used. We thus first inspected obtained patterns visually and found that parameter estimates stabilized after only a few iterations, usually between 10 and 20 (Figure S3).

We then implemented two different measurements to assess convergence more formally: the first is a test statistics assessing the presence of a trend in the parameter estimates, and the second is quantifying the number of slope changes in the individual parameter updates (see Appendix).

### Power to reject Brownian motion

To assess the power of our approach to identify Lévy processes and to estimate associated parameters, we run our EM algorithm on data simulated with jumps on trees of 100 leaves, each simulated using a birth-death model (Stadler, 2011) and scaled to a total length of 1. We generated 100 such simulations for many combinations of number of jumps and *α* values but fixed *μ* = 0 and 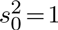 since changing these parameters does not affect the inference. We then inferred the MLE estimates for all parameters under both the null model (Brownian motion) and under the alternative Lévy model.

Using both a likelihood ratio test (LRT) or the Akaike information criterion resulted in generally substantial power to reject the null model over a large range of jumps simulated and for many different values of *α* (Fig. 1). Unsurprisingly, power was much lower if simulated jumps were on the order of the change of the Brownian background process or lower. Here we simulated trees of length 1, and thus the average length of each of the ~200 branches is roughly 0.005. Hence with *α* = 0.01, the strength of half of the evolutionary jumps are expected to be smaller or equal to the effect of the background process on an average branch. However, with *α* = 0.1, the power to reject the null model was > 80% if multiple jumps were present on the tree.

**FIGURE 1.**
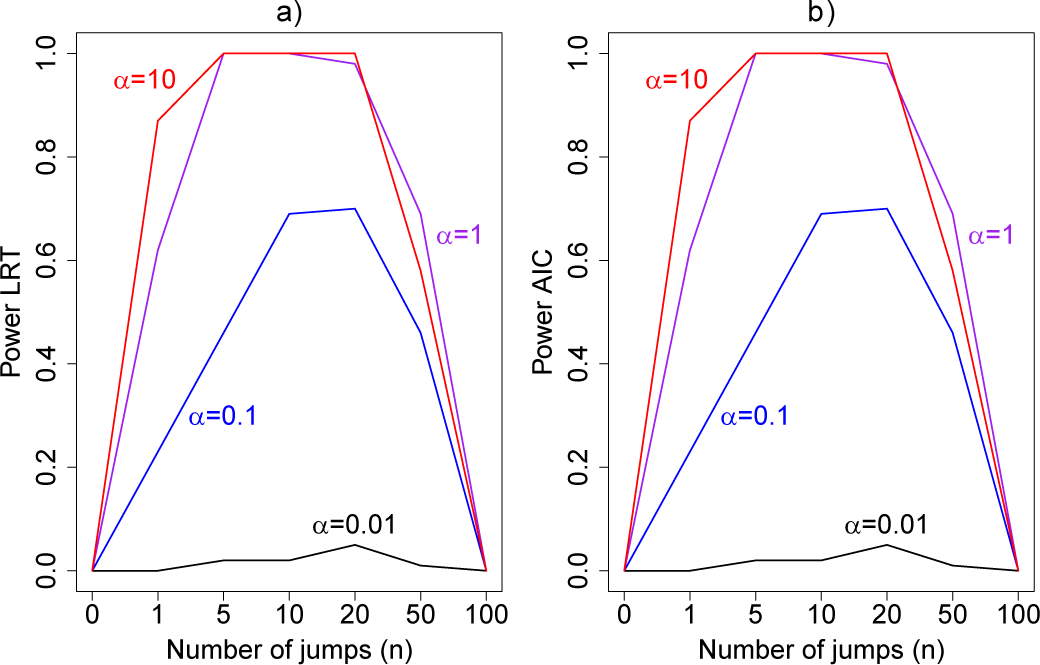
Power to reject the null model (Brownian motion) using a likelihood ratio test (LRT) (**a**), or the Akaike information criterion (AIC) (**b**) as a function of the number of simulated jumps *n* and the jump strengths *α*.

Interestingly, we also found our approach to regularly fail to reject the null model if the number of jumps was very large, i.e. on the order of the number of branches (50 jumps correspond to a jump on every 4th branch). In such situations, the large variance in traits observed under the Lévy model is also perfectly explained by a pure BM model with larger variance 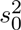 (see below).

In summary, these results show that our method has considerable power to detect Lévy process as long as jumps are meaningfully strong and there are not too many jumps, in which case the Lévy and BM models become indistinguishable from each other.

### Accuracy in inferring Lévy parameters

For the cases in which the Lévy model was preferred we next evaluated the power of our approach to infer the associated parameters, starting with the jump strength *α*. We found that our approach infers *α* quite accurately over the whole range, but we observed a slight overestimation for lower *α* values. This is a direct result of the low power to reject a model of Brownian rate at these lower jump strengths such that for simulations that resulted in larger jumps the Brownian model was more easily rejected. But the inferred values for *α* ≤ 1 were rarely further from the true value than a factor of 2 if multiple jumps were present (Fig. 2a), while it was unsurprisingly much harder to accurately infer the jump strength in case of a single jump.

**FIGURE 2.**
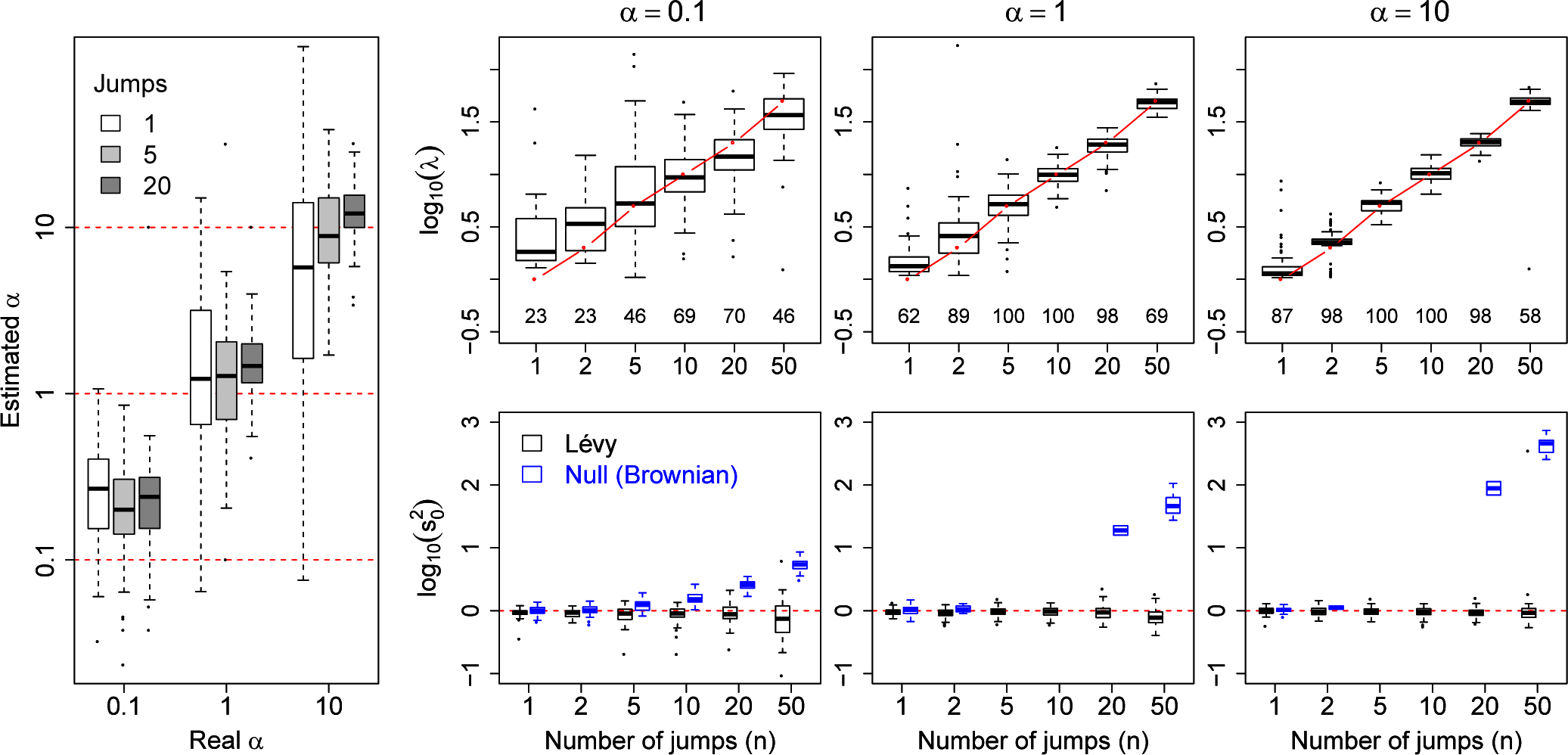
Accuracy in infering Lévy parameters. Each boxplot represents the distribution of inferred values across 100 replicates simulated as described in the text for different combinations of jump strengths *α* and number of simulated jumps *n*. **a)** Accuracy in inferring factor *α*. The true *α* values used in the simulations are indicated with red solid lines. **b)** Top row: distributions of inferred jump rate *λ*. Connected red open circles represent the true values. The numbers printed below the boxplots indicate the percetage of simulations for which the Brownian model was rejected and are hence included here. Bottom row: distributions of inferred Brownian background rates 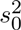 for simulations in which the Brownian null model was rejected (black) or not rejected (blue).

We next evaluated the accuracy of our approach in inferring the jump rate *λ*, again limited to the simulations in which a Lévy model was preferred. As shown in Fig. 2b, our method inferred this parameters very accurately over a large range of jumps simulated and for all values of *α*, with generally higher accuracy with higher *α* values.

We then finally evaluated the accuracy in inferring the Brownian background rate *s*2 (Fig. 2b) and found it to be very accurately inferred whenever the Brownian model was rejected. Interestingly, however, *s*2 was overstimated whenever the Brownian model could not be rejected but jumps were simulated. This illustares that under certain conditions a Lévy model is indistinguishable from a model of pure Brownian motion with an elevated rate. This is particularly true in the case of weak jumps (small *α*) or if jumps are very common on the tree.

### Jump location

We finally tested the power of our method to infer the location of jumps on the tree. For this we simulated trees with 100 leaves and trait data affected by 20 jumps randomly placed on each tree for different jump strengths *α* while fixing *s*2 = 1. In each case we then assumed the Lévy parameters to be known and used our MCMC approach to calculate the posterior probability on there being at least one jump for each branch.

We found our method to have a very low false positive rate in identifying jumps in that a posterior probability for jumps > 0.5 was never obtained for branches on which we did not simulate any jumps (Fig. 3a), and 90% of all such branches resulted in a posterior probability for jumps below 0.2 even for the weakest jump strengths simulated (*α* = 0.1).

**FIGURE 3.**
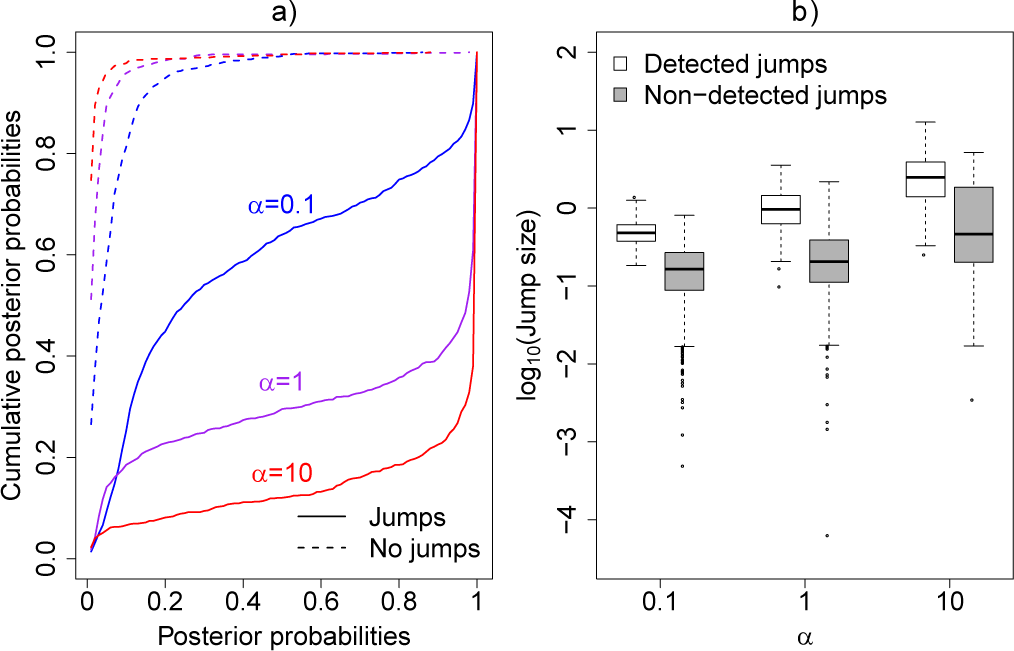
Power to detect individual jumps. **a)** Cumulative distribution of jump posterior probabilities on all branches with (solid lines) and without (dashed lines) simulated jumps. Notice that branches with jumps have posterior probabilities that accumulate at 1, whereas branches without jumps accumulate at 0. **b)** Distribution of the absolute strengths of individual jumps that were either detected (white) or not detected (gray).

The power to infer true jumps (true positives) was also considerably high, especially for jumps of meaningful strength. For data simulated with *α* = 10, for instance, 90% of all branches on which jumps were simulated resulted in a posterior probability > 50%, and 75% even even in a posterior probability > 95%. The few branches with jumps for which we did not obtain decisive posterior probabilities in favor of jumps all contained jumps that were considerably weak (Fig. 3b). Such jumps are expected even for large *α* values since individual jump strengths are assumed to be normally distributed around zero.

A similar pattern was observed when simulating data with smaller *α*, but even in the case of *α* = 0.1 we obtain posterior probabilities in favor of jumps > 0.5 for more than one third of the branches on which jumps were simulated (Fig. 3). At such small *α* values for a tree of length 1, about 40% of all jumps are expected to have a strength smaller than 10 times the effect of the Brownian process on the same branch. But we note that the difficulty in placing weak jumps did not affect the power to infer the jump rate *λ*, which was inferred quite accurately even at such low *α* values (Fig. 2).

## APPLICATIONS

### Quantum evolution in anoles

There have been a few direct tests of Simpsonian jumps between adaptive zones using empirical data (Uyeda et al., 2011). Here, we analyze “evolution by jumps” in the adaptive radiation of anoles, lizards that have adaptively radiated in the Caribbean and South America (Losos, 2009). Following previous work, we focused on anoles on the four islands of the Greater Antilles, as they provide a unique opportunity for testing Simpson's theory of adaptive zones for two reasons. First, there have been repeated dispersal events among islands in the Greater Antilles (Losos et al., 1998; Mahler et al., 2010). These dispersal events represent geographic opportunities, where anole lineages reach a new island and are no longer sympatric with the former set of competitors (Mahler et al., 2010). Second, most anole species can be classified into ecomorphs, habitat specialists that have evolved repeatedly on the four islands of the Greater Antilles (Losos et al., 1998). Transitions between ecomorph categories represent the evolution of key characters in anole lineages that allow them to invade novel habitats (see Losos (2009) for a review).

Anoles have thus repeatedly experienced two conditions under which Simpson expected evolutionary jumps to be observed: dispersal into new geographic areas and the appearance of evolutionary novelties. Importantly, both ecomorph origins and transitions among islands are replicated in the phylogeny of anoles, but are still rare enough that we can estimate the position of transitions on the phylogenetic tree with some confidence (Huelsenbeck et al., 2003; Schluter, 1995).

With this background in mind we tested if a model with evolutionary jumps fits the evolution of body size in anoles better than pure Brownian motion, and if jumps correspond with either of the two factors postulated by Simpson: evolution of key characters and/or geographic dispersal. To address this question, we made use of a recent time-calibrated phylogeny of 170 *Anolis* lizards (Thomas et al., 2009) and analyzed snout-to-vent length (SVL), a standard phenotypic measurement of body size in lizards. This trait is broadly correlated with habitat partitioning in Greater Antillean anoles and represent the primary axes of ecologically driven evolutionary divergence in lizards (Beuttell and Losos, 1999; Losos, 2009; Schoener, 1970). We made use of the sex-specific data of SVL from Thomas et al. (2009) and inferred evolutionary parameters independently for females and males, but excluded five species that lacked information on SVL for one or both sexes (*Anolis darlingtoni*, *A. guamuhaya*, *A. loveridgei*, *A. oporinus*, and *A*. *polyrhachis*).

We found that the Lévy jump model is preferred over a strict BM model in females, but not in males (Table 1). Evolutionary jumps indicating rapid body size evolution (Figure 4) were found precisely at the basis of the clade comprising the ecomorph “crown giants” Thomas et al. (2009), in which females exhibit particularly large body sizes. The large sexual size dimorphism of this group (Harmon et al., 2005) is also likely explaining why the BM model fits the evolution of male body sizes well. In addition to the clades of crown giants, we also identify evolutionary jumps at the basis of the clade consisting of the species *A. barbatus*, *A. porcus*, and *A. chamaeleonides*. These species, which are known as “false chamaleons” and are part of the former genus *Chameleolis* have been called the “most bizarre West Indian lizards” (Leal and Losos, 2000).

**FIGURE 4.**
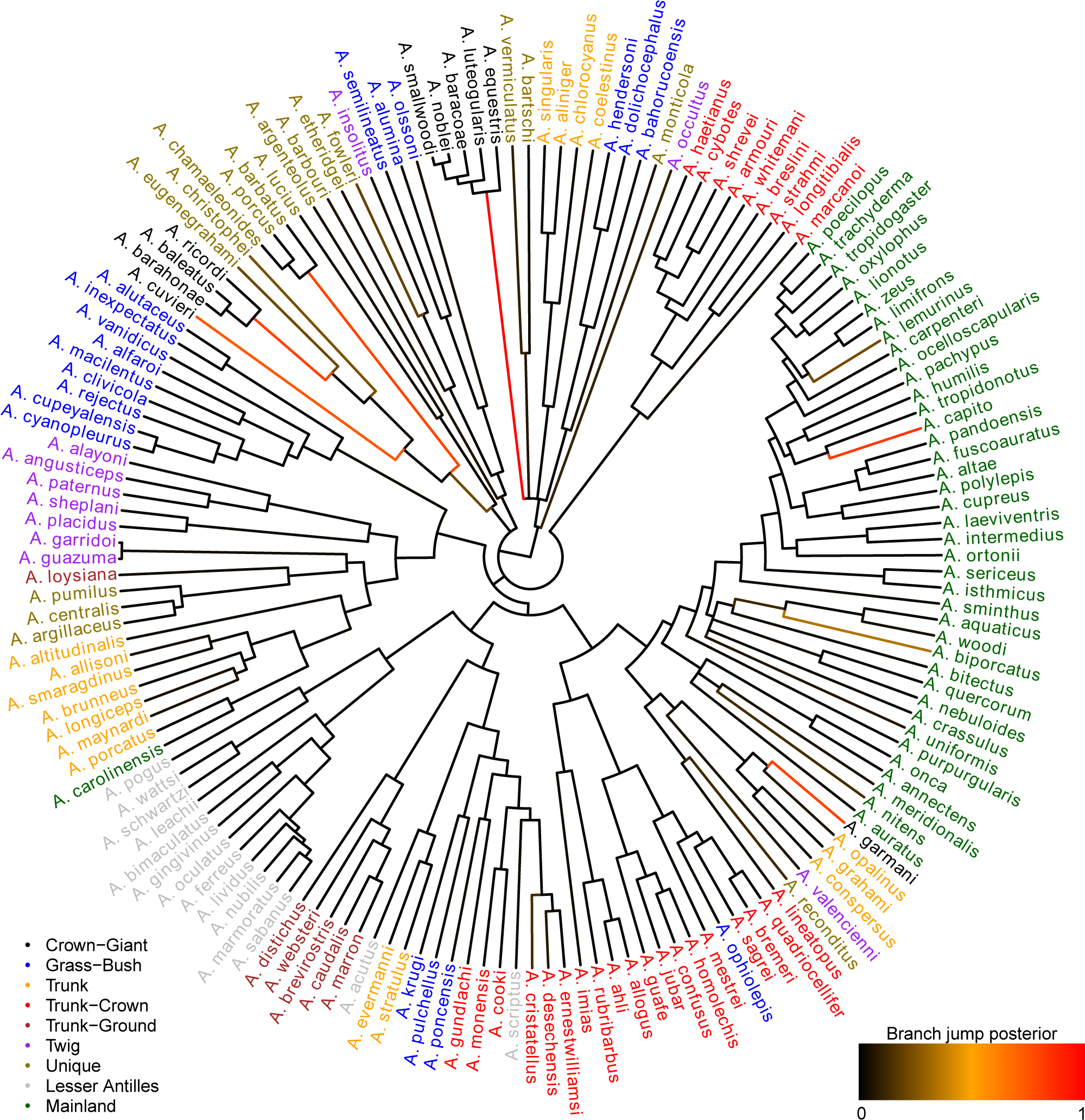
Inferred jumps for female body-size evolution on the anoles' tree. The trait measured was the snout-to-vent length (SVL). Branches are colored according to their inferred jump posterior probability (black to red scale going from posterior probability 0 to 1, respectively). Tips are colored according to ecomorphs as defined by Thomas et al. (2009).

Our analyses support two main conclusions. First, evolutionary change in female anoles is not well described by a uniform random walk. A better description of anole evolution combines a uniform component of change that is punctuated by rapid jumps in trait values. Second, these jumps in body size very well correspond to ecological transitions to novel ecomorphs. The evolution of this trait is thus consistent with Simpson's description of evolutionary jumps associated with the entry into new adaptive zones. The fact that we did not find such jumps at the basis of clades of other ecomorphs suggests that body size was not a trait strongly contributing to the ecological transition of those. However, evolutionary jumps might well be found at the basis of those clades when focusing on more relevant traits.

### Nectarivory evolution in Loriinae

The Australasian lories belong to the tribe Loriini (Schweizer et al. 2015). (Joseph et al., 2012) and are extremely species rich (Schweizer et al., 2011). Their digestive tract is highly adapted to a nectarivorous diet (Güntert, 2012) and Schweizer et al. (2014) has shown quantitatively that a switch in diet to nectarivory might be considered an evolutionary novelty that created an ecological opportunity for species proliferation through allopatric partitioning of the same new niche. Using the methodology developed above we now tested if the evolution of the morphology of the digestive tract in parrots as a whole is better characterized by a model of evolutionary jumps or Brownian motion. For this we made use of data from Schweizer et al. (2014), to generate a time-calibrated phylogeny of 78 parrot species using BEAST (Drummond and Rambaut, 2007) implementing a secondary calibration point from Schweizer et al. (2011) for the initial split within parrots. The following 13 measurements of the digestive tract were used: the length of intestine, length of esophagus, extension of esophagus glands, length of intermediate zone, length of proventriculus, gizzard height, gizzard width, gizzard depth, maximum gizzard height at main muscles, gizzard thickness at main muscles, gizzard lumen width including koilin layer, gizzard width at the caudoventral thin muscle, maximum gizzard height at the thin muscle, and the maximum gizzard lumen at the thin muscle. Since many of the morphological characters of the digestive tract considered are both highly correlated with body size as well as among themselves, we first regressed out body mass (Wgt) from each digestive tract trait and then summarized the residuals of all traits using the first three principal components (PCA; see also (Revell, 2009)).

**TABLE 1.** Inferred Lévy parameters for *Anolis* and Loriini, along with the log-likelihood (*ℓ*) obtained under the Lévy and BM els and the p-value of a likelihood ratio test (LRT) contrasting these.

|  | *Anolis* |  | *Loriini* |  |  |  |
| --- | --- | --- | --- | --- | --- | --- |
|  | logSVL<sub>f</sub> | logSVL<sub>m</sub> | log Wgt | PC1 | PC2 | PC3 |
| $\mu$ | 3.93 | 4.18 | 5.33 | -0.062 | 0.051 | 0.19 |
| $s_0^2$ | 5.06 | 10.34 | 44.40 | 7.68 | 7.35 | 5.80 |
| $\alpha$ | 0.11 | - | - | 0.16 | 0.093 | - |
| $\lambda$ | 11.27 | - | - | 11.95 | 7.38 | - |
| $\ell_{\text{BM}}$ | 5.03 | -15.19 | -85.54 | -65.00 | -60.63 | -6.22 |
| $\ell_{\text{Lévy}}$ | 26.61 | -13.89 | -85.51 | -31.99 | -24.72 | -4.97 |
| LRT $p$ | $4.3 \cdot 10^{-10}$ | 0.28 | 0.97 | $4.7 \cdot 10^{-15}$ | $2.3 \cdot 10^{-16}$ | 0.29 |
| Preferred model | Lévy | BM | BM | Lévy | Lévy | BM |

We found that the evolution of both body weight (Wgt) and the first PC axis of digestive tract morphology were much better explained by a model of evolutionary jumps (*p* < 10^−16^ in both cases) with relatively high rates of jumps (Table 1). Overall, the jumps for PC1 identified with strongest support are both on branches basal to clades of nectarivorous species, particularly at the base of highly specialized nectar feeding Loriini, but also at the base of the genus *Loriculus* (Figure 5). As postulated by Simpson the niche shift to nectarivory especially in Loriini involved a period of rapid evolution reflecting adaptations to feed effectively on nectar (and pollen) (Schweizer et al., 2014). While the jumps within the Neotropical parrots are difficult to interpret in biological terms, the shift along the branch leading to *Psittrichas fulgidus* might be explained by its gizzard morphology similar to that of the Loriini probably reflecting an adaptation to its reportedly mainly frugivorous diet (Schweizer et al., 2014). Some special structures in the digestive tract of the genus *Nestor* have been described in Güntert (2012).

**FIGURE 5.**
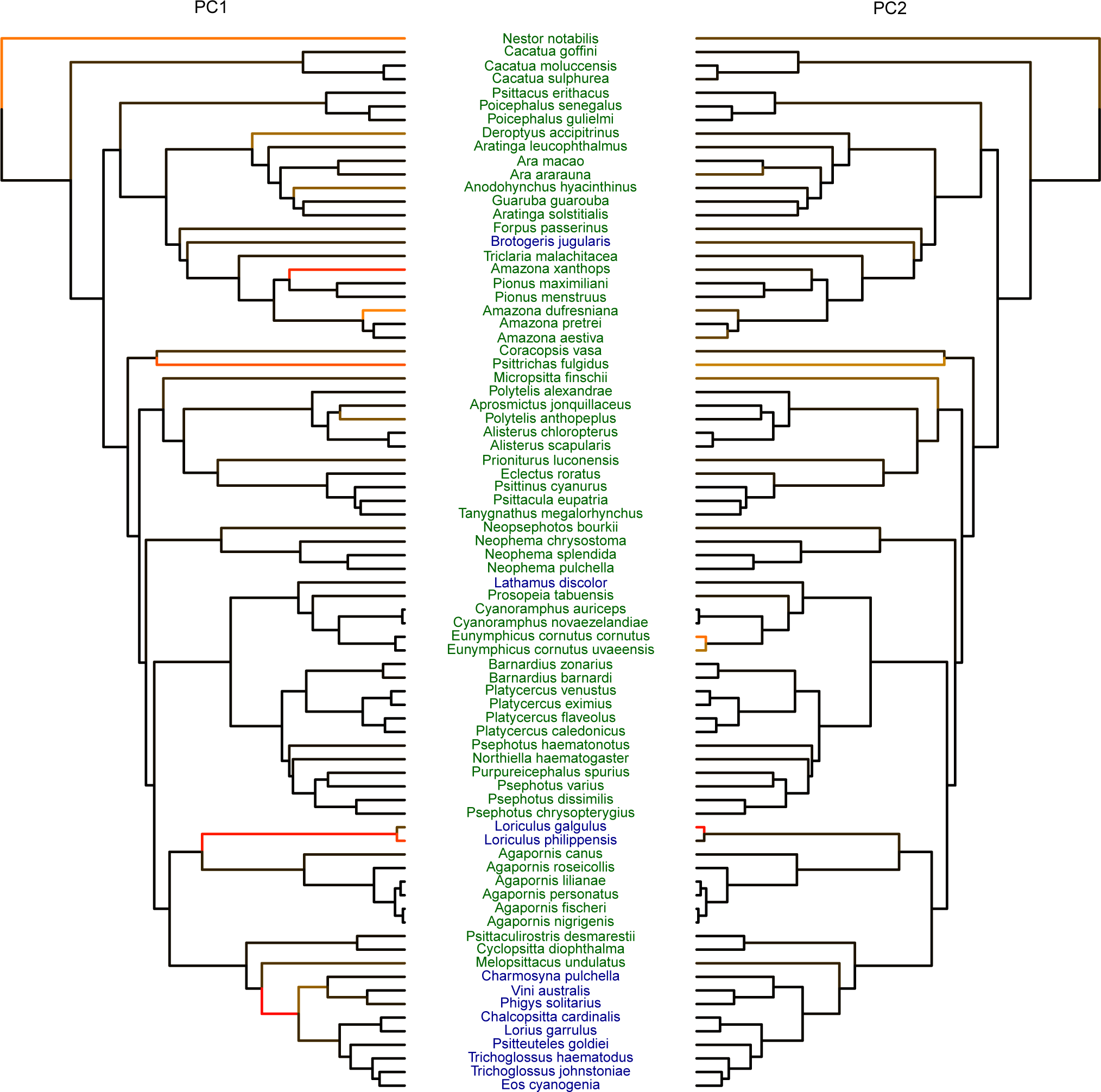
Inferred jumps for I am wondering if we should write more generally here about “evolutionary jumps in the morphology of the digestive tract in parrots”. In any case, it should be Loriini instead of Loriinae. Results for PC1 are shown on the left phylogen and results for PC2 on the right phylogeny. Branches are colored according to their inferred jump posterior probability (black to re scale going from posterior probability 0 to 1, respectively). Species names are colored according to their diet: nectarivorous (blue) an non-nectarivorous (green).

## DISCUSSION

While many traits appear to evolve at relatively constant rates over long time periods and across many taxa, some traits seem to undergo periods of rather rapid evolution (see Arnold, 2014). Simpson (1944) postulated that such evolutionary jumps are triggered by a change in selection pressure after lineages transitioned into different adaptive zones, for instance by dispersing into new geographic areas, after the appearance of evolutionary novelties, key innovations, or after rapid climatic or ecological changes of the environment. The appearance of well calibrated phylogenies along with recent statistical developments now allow to test such models on a wide variety of data.

Bokma (2008), for instance, proposed to model evolutionary jumps as a compound process of a continuous background process and a discrete jump process. Recently, Landis et al. (2013) introduced a general framework to infer parameters of such Lévy processes under a Bayesian framework by means of Markov Chain Monte Carlo (MCMC). Unfortunately this approach, while elegant, requires the calculation of the inverse of the variance-covariance matrix describing the correlations between traits as a function of the phylogenetic tree and the jump process, which is computationally prohibitive for large trees.

Here we introduce a computationally highly efficient variant of this approach that naturally scales to large trees. The basis of our approach is an MCMC algorithm in which we can update the inverse of the above mentioned variance-covariance matrix directly without inversion when sampling jump configurations with fixed hierarchical parameters (root state, Brownian rate, jump strength and jump rate). To make use of this development for inference we propose a two-step approach in which the MCMC algorithm is embedded into an Expectation-Maximization (EM) approach to obtain maximum likelihood (ML) estimates of the hierarchical parameters while integrating over jump configurations. In a second step, the location of jumps can then be inferred under an empirical Bayes framework in which the hierarchical parameters are fixed to their ML estimate and the developed MCMC algorithm is used to obtain for each branch the posterior probability that a jump occurred at this location.

There are also other methods that deal with the burden of calculating inverses and determinants of variance-covariance matrices. For instance, Freckleton (2012) applied the results of Felsenstein (1973) and Felsenstein (1985) to calculate the likelihood in linear time of a BM model. FitzJohn (2012) also proposed a fast algorithm to calculate BM and OU likelihoods using Gaussian elimination, but this is not applicable to non-gaussian traits. Ho and Ané (2014) proposed a new method, which efficiently calculates likelihoods by avoiding the calculation of the inverse and determinant of the variance-covariance matrix. Their method requires that this matrix belongs to a class of generalized 3-point structured matrices. Our method, which applies an iterative scheme, differs from the others in the sense that the inverse and determinant of the variance-covariance matrix has to be calculated only once when obtaining the likelihoods, thus obtaining rather fast calculation times.

We demonstrated the applicability of our approach by identifying evolutionary jumps for body size evolution in *Anolis* lizards and the evolution of the morphology of the digestive tract in Australasian lories of the tribe Lorrini and other parrots. We found strong support for evolutionary jumps in both systems that provide direct support for Simpson's quantum evolutionary hypothesis of adaptive zones. Among the anoles, for instance, we identified evolutionary jumps on the basal lineage leading to crown giants, a group of lizards that transitioned into a novel niche for hunting: the crowns of large tropical trees. Similarly, we identified jumps at the basis of clades of lories that transitioned to nectarivory, an evolutionary novelty that triggered rapid changes in morphology of the digestive system and promoted significant lineage diversification, which was probably mainly non-adaptive after the basal diet shift through allopatric partitioning of the same niche (Schweizer et al., 2014, cf.).

These results also show that the distinction between “gradual” and “punctuated” models of evolution is a false dichotomy; instead, evolution has a gradual component that may be frequently punctuated by periods of rapid change (Levinton, 2001). We further note that in both cases studied here a single jump at the basis of clades is sufficient to explain their trait data, suggesting that the period of rapid evolution was limited to a single branch and that the background rate remained constant. We suggest that future work should follow Simpson's lead and focus on the factors that promote these pulses of evolutionary change.

Although we model evolutionary jumps as instantaneous, we want to be clear that we are not invoking actual instantaneous evolutionary change (e.g. “hopeful monsters”) (Charlesworth et al., 1982; Goldschmidt, 1940). Typical microevolutionary processes of selection and drift can cause change that would appear to be instantaneous when viewed over the timescale of macroevolution. Our model is also distinct from punctuated equilibrium, which requires evolutionary jumps to occur only at speciation events (Eldredge and Gould, 1972). The punctuated changes in our model occur along branches in the tree and are not necessarily associated with speciation events. In fact, for the case of anoles, two lines of evidence argue against punctuated equilibrium: first, most speciation events in the tree are not associated with jumps; and second, we know from detailed microevolutionary studies that anole body size can evolve rapidly in response to selection even in the absence of speciation (e.g. Losos et al. (2006)).

## FUNDING

This study was supported by Swiss National Foundation grant 31003A 149920 to DW. The work of S. M. Szil´agyi was supported by the J´anos Bolyai Fellowship Program of the Hungarian Academy of Sciences.

## APPENDIX

### Conditional likelihood

The EM algorithm can be implemented without the MC part if we impose a condition |***ν***| ≤ *R* on the likelihood, i.e. if we suppose a priori that there have been only *R* or less Poisson events on the tree 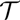. In that case, the sum in (9) is over all *n_k_* such that |*n_k_*| ≤ *R*. Observe that we have to use the conditional probabilities

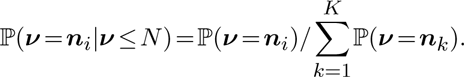

The new *Q*-function is

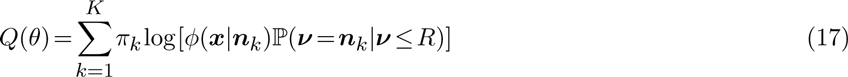

where we can use

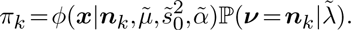

In the conditional case, there no longer seems to exists a closed formula like (12) for the optimal *λ*̃. Setting the derivative of (17) w.r.t. *λ* equal to 0, one can show that *λ*̃ is the root of the following *R*-th order polynomial:

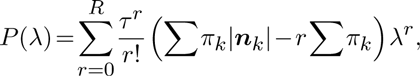

i.e. *P*(*λ*̃) = 0. The estimation of *μ*̃ and *s*̃_0_, on the other hand, remains exactly as explained in section 1.4.

### Assessing convergence of the EM

We introduce two measures to assess convergence of the Monte Carlo EM algorithm.

*Regression criterion* We consider a time series *y*_1_,…,*y*_n_ and construct a test statistic which allows to reject the null hypothesis that the time series exhibits no trend. For this we estimate the slope *β*̂ of the regression line passing through the data points

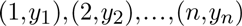

and test for the null hypothesis *β* = 0 (no trend). Determine the following quantities:

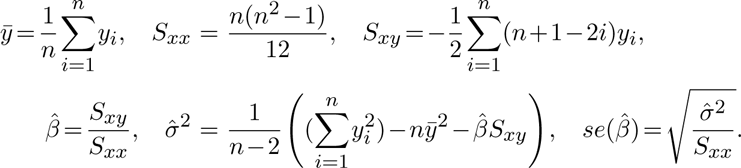

The test statistic

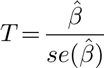

has the Student t-distribution with *n*−2 degrees of freedom. We reject the null hypothesis on the level *γ* if |*T*| ≥ *t_γ_/2,*n*−*2. A good rule of thumb (for *γ* roughly 5% and *n* > 15) is |*T*| ≥ 2.

*Proportion of slope sign changes* We propose a second way of assessing convergence by taking the last *n* values of the EM algorithm and counting the number of times *c* there is a change in the sign of the slope between consecutive values. If convergence is reached, we expect the number of slopes with a positive sign to be similar to the number of slopes with a negative sign. We report the test statistic *N*

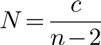

where *n*−2 represents the total number of possible sign changes among the last *n* values.

